# Performance of deep learning restoration methods for the extraction of particle dynamics in noisy microscopy image sequences

**DOI:** 10.1101/2020.11.04.368928

**Authors:** Paul Kefer, Fadil Iqbal, Maelle Locatelli, Josh Lawrimore, Mengdi Zhang, Kerry Bloom, Keith Bonin, Pierre-Alexandre Vidi, Jing Liu

## Abstract

Image-based particle tracking is an essential tool to answer research questions in cell biology and beyond. A major challenge of particle tracking in living systems is that low light exposure is required to avoid phototoxicity and photobleaching. In addition, high-speed imaging used to fully capture particle motion dictates fast image acquisition rates. Short exposure times come at the expense of tracking accuracy. This is generally true for quantitative microscopy approaches and particularly relevant to single molecule tracking where the number of photons emitted from a single chromophore is limited. Image restoration methods based on deep learning dramatically improve the signal-to-noise ratio in low-exposure datasets. However, it is not clear whether images generated by these methods yield accurate quantitative measurements such as diffusion parameters in (single) particle tracking experiments. Here, we evaluate the performance of two popular deep learning denoising software packages for particle tracking, using synthetic datasets and movies of diffusing chromatin as biological examples. With synthetic data, both supervised and unsupervised deep learning restored particle motions with high accuracy in two-dimensional datasets, whereas artifacts were introduced by the denoisers in 3D datasets. Experimentally, we found that, while both supervised and unsupervised approaches improved the number of trackable particles and tracking accuracy, supervised learning generally outperformed the unsupervised approach, as expected. We also highlight that with extremely noisy image sequences, deep learning algorithms produce deceiving artifacts, which underscores the need to carefully evaluate the results. Finally, we address the challenge of selecting hyper-parameters to train convolutional neural networks by implementing a frugal Bayesian optimizer that rapidly explores multidimensional parameter spaces, identifying networks yielding optional particle tracking accuracy. Our study provides quantitative outcome measures of image restoration using deep learning. We anticipate broad application of the approaches presented here to critically evaluate artificial intelligence solutions for quantitative microscopy.

## INTRODUCTION

Sample illumination is a conundrum in live cell imaging: too much light leads to fluorophore photobleaching and phototoxicity (Carlton *et al.*, 2010; Magidson and Khodjakov, 2013), whereas too little light results in poor images with low signal-to-noise ratio (SNR). Low SNR is detrimental to quantitative microscopy applications requiring precise localization and tracking of particles (Chenouard *et al.*, 2014). The issue of illumination vs. SNR is compounded in single molecule tracking (SMT) experiments because each fluorescent molecule emits a limited number of total photons that can be collected (so-called photon budget). In addition, fast imaging is often needed to precisely map particle trajectories, in particular for those particles that move rapidly. High-speed imaging requires short exposure times which in turn gives low SNR.

The different types of noise and the relationship between exposure time and SNR for cameras (e.g. CCD and sCMOS) are described by the equation (Eq.1; (Fellers and Davidson, 2004) and the schematic in Box 1. Shot noise is due to the statistical nature of the rate of photons arriving on the camera, and the interval between arrival of photons is defined by Poisson statistics. This noise is captured by the factor (*S*+*B*) in the denominator of Eq.1. An additional source of noise included in Eq. 1 is the background *B* generated by radiating fluorophores above and below the imaging plane, by scattered light in the microscope, as well as by autofluorescing objects in the sample. Dark noise is due to the statistical variation of the electrons thermally generated by the camera. For a given camera, we assume constant *P*, *QE*, *N_r_* and *D_c_*. Dark noise is negligible for exposure times of 1 s or shorter compared to the typical read noise at standard frame rates of *N_r_* = 1.4 electrons root mean square (rms). Read noise is the noise generated by the amplifying electronics in a camera that converts the electrons generated at a pixel into a voltage output and then into a digital value via an analog-to-digital converter. A key difference between CCD and sCMOS cameras is that the read noise is almost identical for each CCD pixel, whereas sCMOS pixels have read noise that is pixel-dependent. Note that read noise is squared in its contribution to the noise denominator in Eq. 1. Therefore only the signal and photon shot noise play significant roles in the variation of SNR with exposure time *τ*.

Recognizing the various sources of noise and classifying their statistical features is at the core of several classic noise reduction algorithms. These include BM3D (Danielyan *et al.*, 2012) and nd-safir (Boulanger *et al.*, 2010). Both methods are based on collaborative filtering: they group together image patches with similar statistical properties and apply transformations that use all grouped patches to restore each grouped patch. The methods differ in how they define patch similarity, as well as in how they use the groups to restore patches. Nd-safir restores a patch by using the weighted average of the group, with more similar patches having greater weight in the average. BM3D weighs patch-contributions within the group by its estimate of the noise they contain: less noisy patches carry greater weight. While effective in increasing the SNR of input images, these approaches are content agnostic: they use the statistical properties of the intensities within the input images, without regard to the image content. This makes them applicable to any dataset as long as their assumptions are met. However, in some cases it may be beneficial to sacrifice generalizability and use approaches that are dataset-specific to achieve a further increase in the SNR and superior restoration. Denoising approaches that make this trade-off are called content-aware: they collect and use information about the content of the dataset, and can achieve higher SNR and notable qualitative image improvements as a result of exploiting this additional information.

With the rapid improvement in GPUs, deep learning has revolutionised quantitative image analysis problems, including cell segmentation, object classification, and particle counting and tracking (Moen *et al.*, 2019). In addition, deep learning methods that use empirical knowledge on noise have a great potential to restore low-SNR datasets, as mentioned above. The content-aware image restoration (CARE) network developed by Weigert et al. (Weigert *et al.*, 2018) and based on the the U-net convolutional neural network (CNN) architecture (Ronneberger *et al.*, 2015) is a popular tool for image restoration. In its original implementation, CARE uses pairs of images with high and low noise levels to generate data-specific denoising networks. High-quality ‘ground truth’ (GT) images with high SNR are not always available, and self-supervised approaches have been developed that eliminate the need for GT. One example is Noise2void (N2V) (Krull *et al.*, 2018, 2020). The innovation that allows N2V to denoise images with only access to a single noisy image (or set of images) is the introduction of blind-spot networks. The N2V network is given an image patch, and tasked with predicting the center pixel’s value. A conventional U-net would learn to directly output the center pixel’s value, ultimately leaving the input image unchanged. In contrast, the N2V network has a blind spot at the center pixel, forcing it to infer the center pixel’s intensity from the surroundings. Krull et. al. have shown that such blind-spot networks learn to remove pixel-wise independent noise.

Image restoration with deep learning for quantitative microscopy is still in its infancy. While image improvement with CARE and N2V (among other deep learning approaches) is qualitatively impressive, to the best of our knowledge, the ability of these neural networks to restore data for quantitative analyses of molecular properties, such as diffusive behaviors, has not been rigorously evaluated. Particle tracking is an ideal test case to objectively assess the performance of image denoising approaches; specifically, their ability to restore meaningful biological information, beyond cosmetic improvements. Here, we compare tracking performances after denoising with supervised and unsupervised CNNs, using synthetic image sequences of diffusing beads as well as chromatin dynamics as a cell-biological application.

## RESULTS AND DISCUSSION

### Restoring images of diffusing particles: proof-of-concept with synthetic movies

As an initial step to test if CNN-based image denoising can improve accuracy in particle tracking experiments, we generated synthetic time-lapses of moving beads. With this approach, the number of particles, their diffusion coefficient (*D*), background shot noise, and readout noise were defined for each movie. In these artificial movies, all particles underwent Brownian motion and their mean squared displacement (MSD) was used to describe changes in the positions of the particles with respect to the characteristic time interval (*τ*). For denoising with CARE, pairs of movies were created that had either no added noise (ground truth) or different levels of added noise (Fig. 1A and Suppl. Fig. S1A). A subset of the data was used to train CARE networks, which were then applied to denoise the remaining noisy movies. N2V only requires a series of noisy images for training, without corresponding GT. The beads in both denoised and GT movies were detected and tracked using a single-particle tracking algorithm with cross-linking to develop the trajectories (see Materials and Methods).

**Figure 1.**
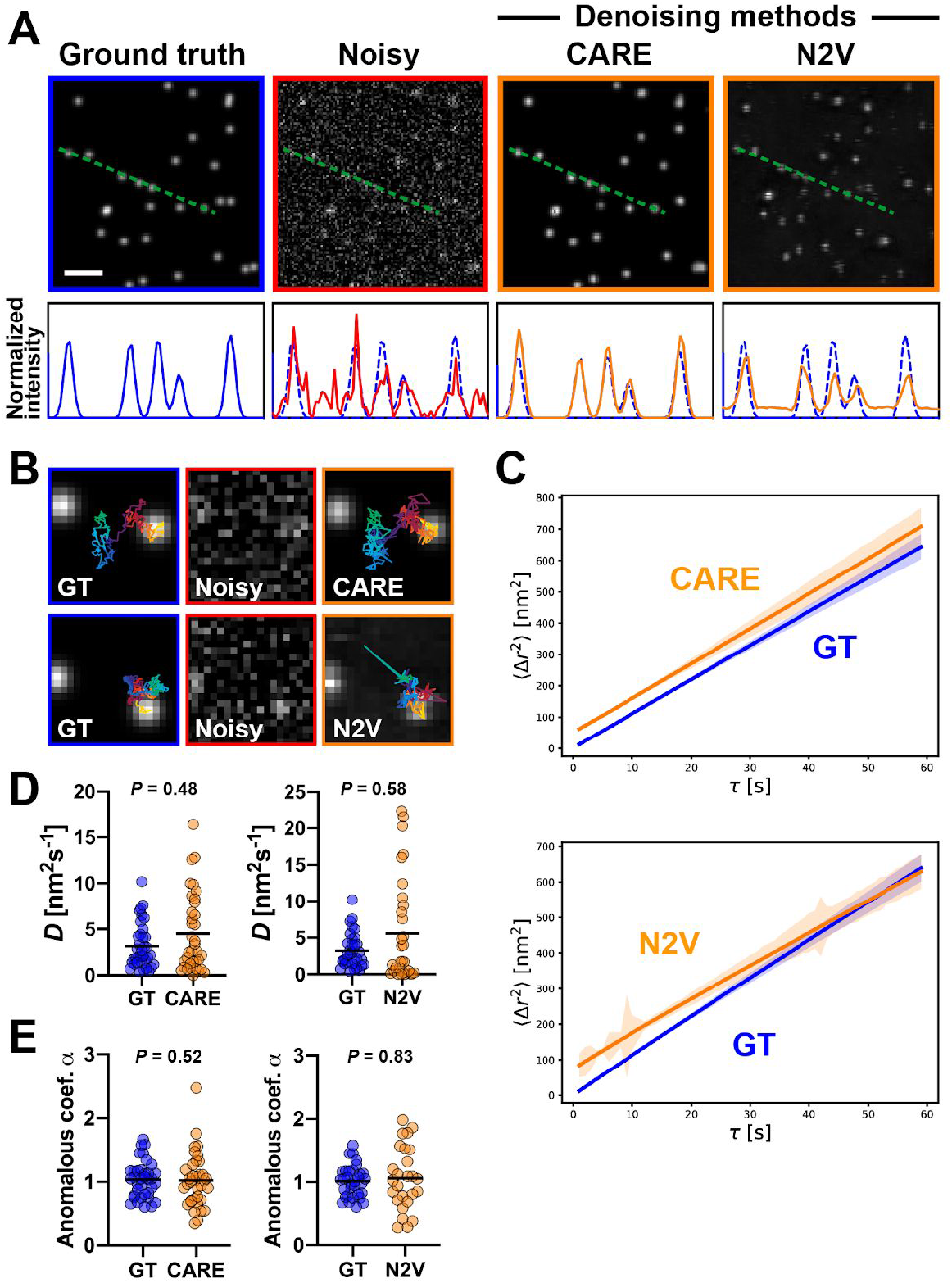
Evaluation with synthetic data of the performance of supervised and unsupervised CNN denoising approaches for particle tracking. **A** Representative synthetic images of diffusing beads, without added noise (ground truth) or with noise added (*σ* = 0.5, see Materials and Methods). Noisy images were denoised with different algorithms, as indicated. Intensity profiles along the dotted lines are shown. Scale bar, 2 μm. **B** Illustration of particles trajectories in ground truth (GT) and denoised movies. Particles in noisy movies could not be tracked. **C** Averaged MSD curves from beads tracked in ground truth and denoised movies. **D** Diffusion values computed from the MSD curves. **E** Anomalous diffusion coefficient (*α*) calculated from denoised and GT MSD curves. Statistical comparisons (D and E) using Mann Whithney test. Mean values are indicated.

As expected, both CARE and N2V improved noisy synthetic movies, revealing beads barely distinguishable by the human eye in images with high noise levels. This improvement can be appreciated with the intensity profiles shown in Fig. 1A. Beads in movies with relatively low noise levels (σ = 0.2) could be precisely tracked after denoising (Suppl. Fig. S1B-C). In movies with high noise levels (σ = 0.5), for which tracking was not possible, CARE denoising enabled correct identification of ∼75% of the particles (Suppl. Fig. S1D). Qualitatively, movies denoised with CARE showed individual particle trajectories closely resembling the ones extracted from the ground truth. In contrast, N2V denoising lead to more dissimilar trajectories (Fig. 1B). Quantitatively, MSD curves derived from movies processed with the CARE algorithm coincided with the ground truth (Fig. 1C and Suppl. Fig. S1C). Accordingly, diffusion coefficients and anomalous exponents derived from the denoised movies were consistent with the theoretically defined values and not different from the ground truth (Fig. 1D-E). Similarly, untrackable movies could be analyzed after denoising with N2V, although tracking errors occurred more frequently compared to CARE. To further assess the accuracy of the denoised movies, we calculated the least distances between bead spot centers in GT and denoised movies. The cumulative values for these tracking errors are shown in Suppl. Fig. S1F. After denoising with the CNNs (CARE and N2V), the mean cumulative localization error (over 300 frames) was ∼50 nm (or ⅕ of the size of the point spread function). Localization error was slightly greater for bead image sequences denoised with N2V. It should be noted that this localization error did not deteriorate the restored particle trajectories; it just shifted the baseline of the MSD curves without affecting fitting results of *D* values (Fig. 1C).

In summary, these experiments with synthetic images of diffusing particles show that CNN-based image restoration has the potential to not only improve the image quality, but also to recover the motion behavior of moving particles. We find that supervised CARE networks can restore the dynamics of single beads with greater accuracy than unsupervised N2V.

### Content-aware denoising for tracking chromatin microdomains in live cells

After establishing proof-of-concept with a synthetic dataset, we applied the CARE and N2V softwares to noisy time-lapses of slow-diffusing chromatin microdomains to evaluate if and to what extent CNN-based denoising can restore kinetic information in a biological dataset. We used a custom optical setup based on diffractive optics to photoactivate 7×7 lattices of chromatin microdomains in U2OS cells stably expressing histone H2A tagged with photoactivatable GFP (PAGFP-H2A) (Bonin *et al.*, 2018) (Fig. 2A). Interlaced movies were collected from the photoactivated lattices, alternating short and longer exposure times. Because the time difference between adjacent frames (short - long exposures) is small, we consider the image sequence taken with long exposures as the ground truth for the noisy (N) image sequence. We applied the same approach with different short exposures (10 ms, 3 ms, and 1 ms), corresponding to increasingly noisy images. To restore noisy image sequences, we used CARE networks trained on N/GT image pairs. Alternatively, we trained N2V networks tailored to individual noisy sequences with the goal to maximize the outcome of unsupervised denoising.

**Figure 2.**
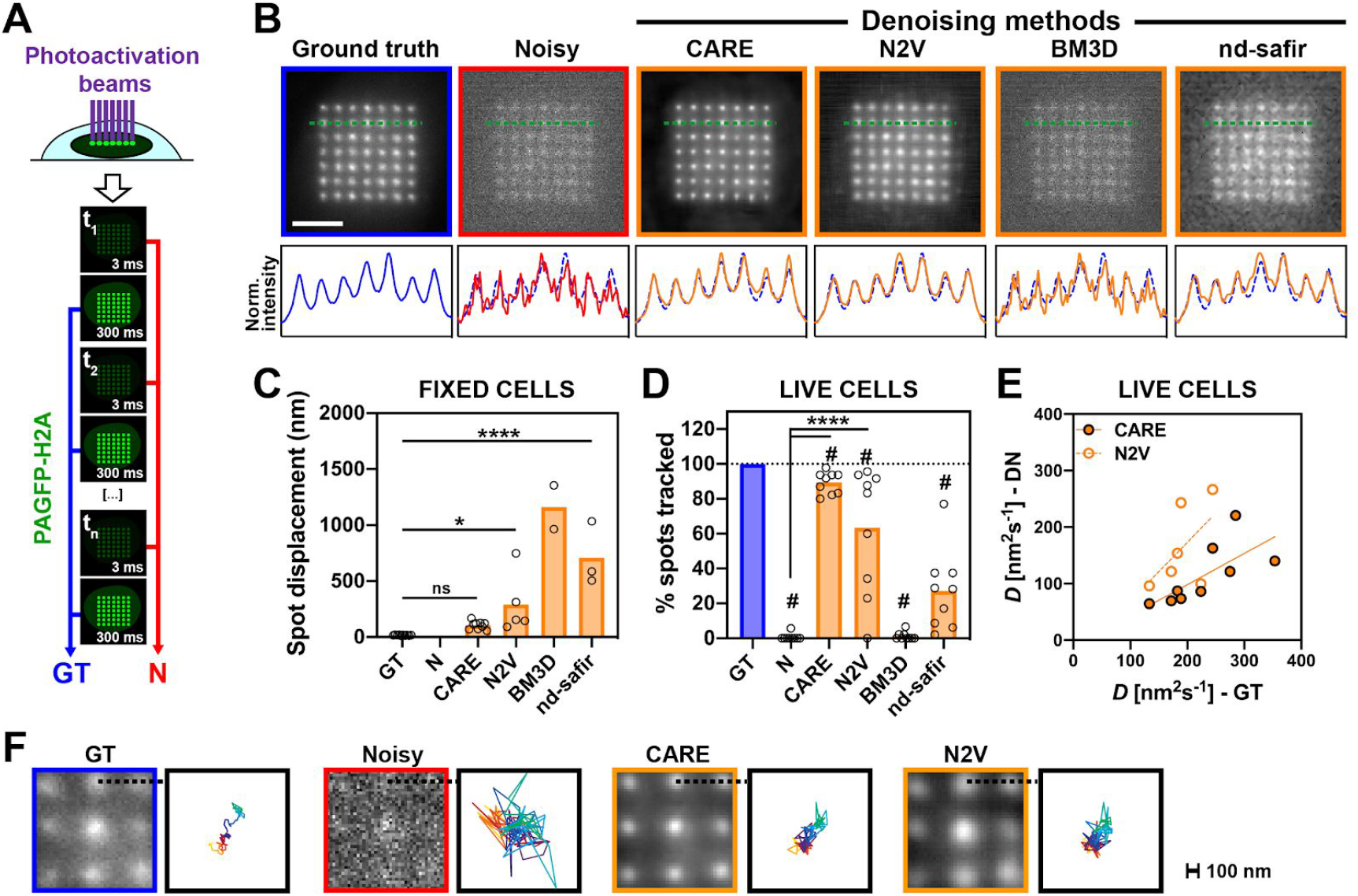
Image denoising to track chromatin microdomains in live cell nuclei. **A** Schematic of the approach to photoactivate chromatin microdomains in U2OS cells expressing PAGFP-H2A. Rapid successions of short and longer exposure times were collected. **B** Representative images of photoactivated chromatin microdomain lattices taken with a long exposure (ground truth), with a short exposure (noisy), and after denoising with different algorithms. Pixel intensity profiles are shown across a row of microdomains. Scale bar, 5 μm. **C** Cumulative spot motions in fixed cells. No microdomain could be tracked in the noisy dataset. **D** Percentages of photoactivated spots that could be tracked in noisy movies or after denoising with the different algorithms. For each cell, values are normalized to the number of spots tracked in the GT. **E** Chromatin diffusion (*D*) in live cells, comparing GT and denoised (DN) movies. Each dot in the graphs (D-E) represents the average value for a cell. **F** Representative traces of microdomain trajectories. *, *P* < 0.05; ****, *P* < 0.0001; ns, not significant (ANOVA and Tukey’s test). #, *P* < 0.05 (one sample t-test; theoretical mean of 100). Mean values are indicated in the bar graphs.

Qualitatively, images denoised (DN) with CARE or N2V were much sharper than the original noisy images and resembled the GT (Fig. 2B and Supl. Fig. 2A). Accordingly, pixel intensity profiles across photoactivated spots revealed clear peaks that appeared to be similar in GT and DN images. Even for movies with the highest noise levels (1 ms exposure), improvement of the noisy images with CARE was striking at first sight (Supl. Fig. 2B). Yet, closer examination revealed striking differences in the DN outcomes, when comparing images restored from very low (1 ms) vs. low (10 ms) exposure data. In DN images from very low exposures, fluctuations of background between frames were exacerbated, spot shapes appeared to have reduced complexity, and - most concerning - DN images occasionally included hallucinated spots inside and outside of the photoactivated lattice. Hence, denoising results need to be carefully evaluated in situations with extreme noise levels, in particular when repetitive elements are present in the images.

To compare tracking accuracies achieved with GT, noisy, and DN data, image sequences were registered to cancel cell motions and the center position of each chromatin microdomain in each image frame was determined by fitting with a 2D Gaussian function. We used these positions to calculate MSD curves and diffusion coefficient *D* values (Bonin *et al.*, 2018). In fixed cells, we expect no microdomain motion; residual diffusion reflects drift of the microscope stage that was not properly subtracted, combined with imprecise Gaussian fitting leading to errors in spot center positions. As a first step to compare the DN methods, we calculated cumulative spot displacements in fixed cell movies (Fig. 2C and Supl. Fig. 2C). As expected (Bonin *et al.*, 2018), spots in GT movies from fixed cells barely moved (*D* = 2.2 ± 1.3 nm^2^/s). In contrast, the apparent spot motions in the corresponding noisy movies (10 ms exposure) were ten times greater, and most spots from movies with greater noise levels (3 ms exposure) could not be tracked, partly due to failure in image registration (Supl. Fig. S2F). CARE and N2V denoising reduced the apparent spot motion of noisy images. Increased accuracy in spot center determination and improved image registration both contributed to this improvement in the DN movies (Supl. Fig. S2E). Next, we used the same approach with live cells. The proportion of chromatin microdomains that could be tracked significantly increased after denoising with CARE (Fig. 2D and Supl. Fig. S2D). Denoising with N2V also increased the proportion of trackable spots, albeit to a lesser extent. To assess denoising outcomes, *D* values from DN movies were plotted against the corresponding values in GT (Fig. 2E). Restoration of 3 ms movies to the 300 ms GT with CARE led to DN-derived diffusion values that were significantly correlated with GT-derived *D* values (r = 0.72, *P* = 0.029). Moreover, individual microdomain traces from DN spots were qualitatively more similar to GT than to noisy source images (Fig. 2F). Although CNN-based denoising (and CARE in particular) enabled tracking of >80% of the microdomains and yielded diffusion values correlated with those from GT, we note that *D* values in CARE-DN movies were systematically lower than their GT. This effect may be due to the loss of dynamic changes in spot shape after denoising. Microdomain spots in our experiments have a typical size of 600 nm and nonsymmetric shapes are usually observed in the GT, whereas images denoised with CARE display more symmetric spots with Gaussian profiles. It is likely that fluctuating structural variations are interpreted as noise by the software. Another limitation in our comparisons is that the unsupervised CNN that we used (N2V) cannot identify nor remove structural noise (apparent as horizontal and vertical stripes in the N2V-DN images). It will therefore be interesting to assess the performance of StructuredN2V (Broaddus *et al.*, 2020), a generalized version of N2V using blind-spot masks based on noise structure.

To benchmark the results obtained with CNN-based denoising, the same noisy image sequences were processed using two software solutions not based on deep learning, BM3D (Danielyan *et al.*, 2012) and nd-safir (Inria©) (Boulanger *et al.*, 2010). Both qualitatively and quantitatively, BM3D did not achieve CNN-based denoising outcomes (Fig. 2B-D, Supl. Fig. S2A,C,D). This was expected since this software is best suited for images with a large proportion of signal rather than dominance of background pixels. In addition, BM3D requires an estimate of the noise variance, which was not trivial to assess. Overall, results obtained with nd-safir with moderately noisy images (10 ms exposure) were comparable to those obtained with unsupervised deep learning (Supl. Fig. S2). With higher noise levels (3 ms exposure), movies denoised with nd-safir could not be tracked (Fig. 2). The nd-safir software can handle 4D datasets. We may therefore have underestimated the performance of this software to restore tracking information since we did not exploit the temporal component of the movies. Nevertheless, the results suggest that CNN-based denoising outperforms classic approaches, at least for this specific quantitative microscopy application.

### Application of CARE for tracking single molecules in live cells

Next, we applied CNN-based denoising to single-molecule imaging by tracking the dynamics of individual nucleosomes in nuclei with stochastically-labelled histone H2B. For these experiments, we used U2OS cells expressing H2B fused to the HaloTag, to which a fluorescent ligand can bind specifically (Liu *et al.*, 2018). Stochastic labeling of H2B was achieved by incubating U2OS H2B-HaloTag cells with a low concentration of fluorescent ligand, for a short amount of time. For imaging, live cells were illuminated by an oblique light sheet and the fluorescent signal was collected by a camera at a rate of up to 100 frames/second (Fig. 3A). Compared to microdomains of chromatin, single nucleosomes have much faster kinetics that can only be captured by imaging at high frame rates (Nozaki *et al.*, 2017). And similarly to other SMT experiments, the photon budget of each labeled nucleosome is limited, meaning that low illumination intensity is needed to capture time-lapse series of meaningful length. To train CNN denoising networks, we used fixed cells and captured matching pairs of low exposure (10 ms or 50 ms) and ground truth (1 s) images. For CARE, both noisy and ground truth images were used for supervised learning, while only the noisy data were used for N2V training. We used the same approach as the one used for chromatin microdomains, recording movies alternating short (10 ms) and longer (50 ms) exposures (Fig. 3A-B). The trained CARE and N2V networks were applied on the short exposure movies, while the longer exposure movies were used as ground truth.

**Figure 3.**
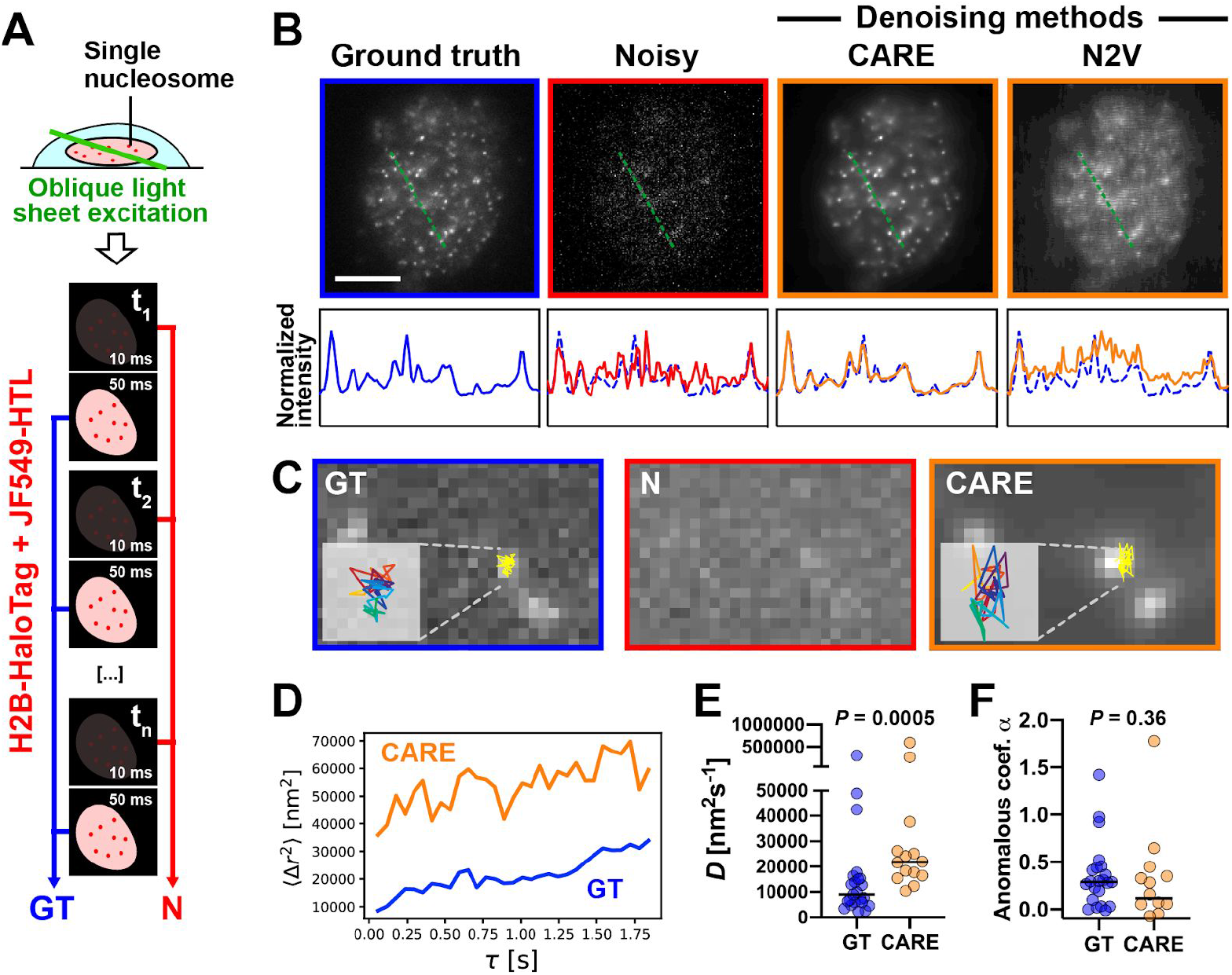
Restoration of the fast dynamics of nucleosomes captured with single-molecule imaging. **A** Schematic of the oblique light sheet imaging approach with U2OS cells expressing H2B-HaloTag. Rapid successions of short (10 ms) and longer (50 ms) exposure times were collected, leading to matching noisy (N) and ground truth (GT) movies. **B** Representative ground truth and noisy images, as well as images denoised with CNN algorithms. Pixel intensity profiles along the dotted line are shown. Scale bar, 10 μm. **C** Illustration of particles trajectories in GT and denoised (CARE) movies. Particles could not be tracked in noisy movies. **D** Representative MSD curves. **E** Diffusion values (*D*) of single nucleosomes in GT and denoised movies. **F** Comparison of anomalous diffusion coefficients (*α*) derived from single nucleosome MSD curves. Statistical comparisons (E and F) using Mann-Whitney test. Median values are indicated.

Again, the performances of CARE exceeded that of N2V in restoring single molecule images, as indicated by the intensity profiles of individual nucleosomes (Fig. 3B). CARE restored both the localization and relative fluorescence intensity of nucleosome foci, whereas N2V only partially recovered the bright spots from the raw image, with “scanning-like” patterns for each foci, and intensity profiles indicating a significant mismatch with the ground truth. Therefore, it is clear that the supervised denoising approach is better suited than the unsupervised approach for single-molecule image series, as the pattern and statistical features of the noise in these datasets are largely unknown.

Next, we evaluated the dynamics of the nucleosomes following CNN-based image restoration. We had established that movies of nucleosomes taken with a 50 ms exposure can be used for single-molecule tracking with our system, whereas 10 ms movies could not be tracked due to the high background noise. First, we verified if CARE denoising of ‘trackable’ nucleosome movies induces artifacts. We obtained similar tracking results with 50 ms exposure images and the same images processed with the CARE network described above (Supl. Fig. S3). We then focused on high-noise movies. As illustrated in Fig. 3C, trajectories of the same nucleosome tracked in both 50 ms exposure (GT) and denoised 10 ms exposure movies were qualitatively similar. More importantly, the calculated MSD curves based on the GT trajectories had slopes and shapes matching the ones derived from the 10 ms denoised trajectories (Fig. 3D), with the y-axis shift due to the larger localization error of DN spots. Diffusion values derived from the denoised movies were about two-fold larger than the GT movie. This difference can be explained by the fact that motion blur is greater for longer exposure movies and leads to an underestimation of particle velocity. Statistically, the α values were similar for the GT and DN movies (Fig. 3F) and lower than 0.5, indicating that the subdiffusive behavior of nucleosomes (Nagashima *et al.*, 2019) was captured in DN movies.

### Denoising and tracking of three-dimensional (3D) dataset

Results presented so far were all derived from simulated or experimental two-dimensional images, with positions and trajectories projected and extracted from a single z-plane. While these types of measurements are practical and generally well-suited for flat samples, such as cell nuclei in monolayer cell cultures, 3D measurements improve particle tracking accuracy, in particular for round nuclei where the relationship between 2D vs 3D distances deteriorates, and for short distances (< 5 μm) where the average 2D/3D discrepancy is ∼30% (Finn *et al.*, 2017). Tracking of genomic loci in the small, round, and tumbling nuclei of yeasts is one instance where 3D datasets are required (Gasser, 2002; Chacón *et al.*, 2014; Lawrimore *et al.*, 2018). To assess particle tracking performance after CNN-based denoising of image volumes, we generated simulated 3D microscopy time-lapses of the budding yeast pericentric region (Lawrimore *et al.*, 2016) with different noise levels (See Methods section). These movies model DNA loops containing 200 bp or 400 bp fluorescent reporter operator arrays (Fig. 4A). The loops exhibit confined motion due to the tethering of the simulated DNA at the centromere. The 400 bp array signal is twice as bright as the 200 bp array signal. Fig. 4B shows representative images of the simulated loci, with the different noise levels. Image sequences without any added noise are shown as ground truth comparison.

**Figure 4.**
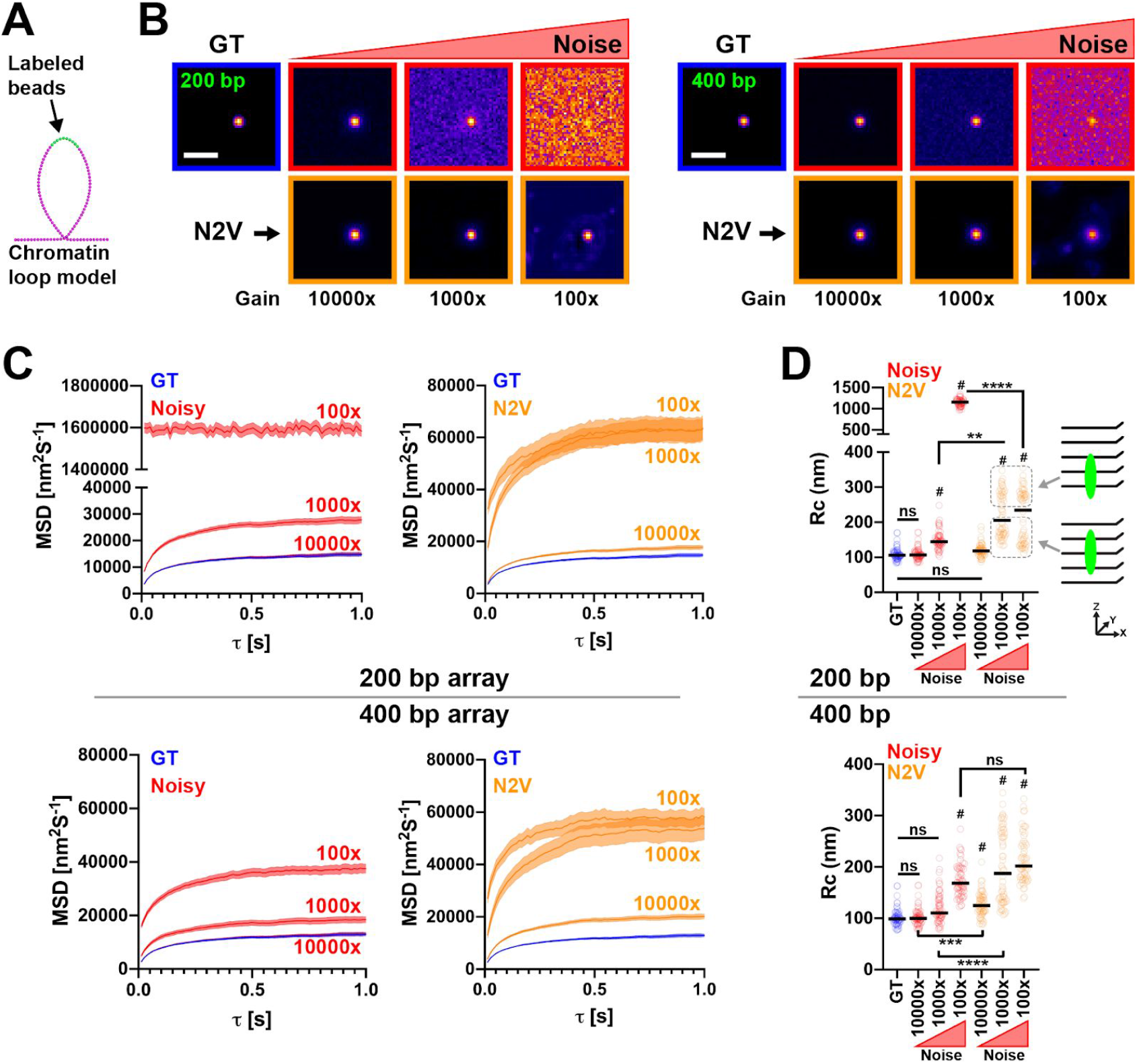
Effect of CNN-based image volume restoration on tracking performance for single chromosomal loci. **A** Schematic of the fluorescent labeling of a single loop from a ChromoShake simulation of the budding yeast pericentromere (Lawrimore *et al.*, 2016). **B** Representative images generated from ChromoShake simulations with 0 noise (GT) and normally distributed random noise with a standard deviation of 5 AU. The noised images were generated with 100-fold, 1,000-fold, and 10,000-fold gains to create images with differing signal-to-noise ratios. Each image is displayed to show the image’s full intensity range. The contrast differs between images. **C** Ensemble MSD curves of GT and denoised image foci of the simulated 200 bp (top) and 400 bp (bottom) arrays. Each plot is composed of 64 different loops from a single pericentromere simulation. Error bars represent standard error of the mean. Initial localization of the focus within the noised 100x images of 200 bp array and of 400 bp array exceeded the cropping region of the 3D Gaussian fitting in 23.45% and 0.004% of images, respectively (see Methods). The large percentage of mis-localized foci in the 100x images of the 200 bp array caused the large Rc values in D. Ground truth MSDs were calculated directly from simulation coordinates. **D** Comparison of radii of confinement (Rc) from GT and tracked images for 200 bp and 400 bp arrays. The schematic shows that the higher Rc values correspond to foci that are lower in the z-dimension for the denoised images of the 200 bp array. **, *P* < 0.001; ***, *P* < 0.0005 ****, *P* < 0.0001; #, *P* < 0.0001 compared to GT; ns, not significant (Kruskal-Wallis and Dunn’s multiple comparison test). N = 64 simulated time-lapses. Median values are indicated.

The content-aware denoising software that we have so far evaluated can also be trained on 3D data. Since ground truth images are rarely available for existing biological datasets, and considering that N2V is fully implemented in ImageJ, an image analysis software broadly used in the cell biology community, we focused on unsupervised denoising with N2V. The simulated hyperstacks were denoised using N2V, with one N2V network trained for each imaging condition on all images of that condition. The foci were tracked using a Gaussian fitting approach and the MSDs at different time intervals were calculated for each of the tracks (Fig. 4C). The plateau value of each MSD curve was converted into a radius of confinement (Rc), which represents the area a focus can explore. The ground truth MSDs and confinement radii were calculated directly from simulation coordinates and compared to values from noisy and N2V-denoised simulation movies. As expected, time-lapses with low noise levels could be tracked precisely, yielding Rc values not statistically different from their respective ground truth values (Fig. 4D and Table S1). For the dimmest array (200 bp), processing with N2V did not significantly alter the tracking outcome (Fig. 4D-C, compare 10,000 gain and GT). For the other brightness and noise conditions, denoising with N2V leads to significant overestimation of the Rc values, compared to both GT and noisy input movies. Only in the most extreme condition (dimmer 200 bp array and highest noise level) did N2V denoising significantly improve tracking outcome. For these images, in which individual loci were barely distinguishable and tracking essentially failed (Fig. 4D top, 100x gain), N2V denoising enabled tracking, yielding Rc values overestimated by a factor two. The bimodal distributions present in the denoised 200 bp array images are due to significantly more variation in the z-dimension tracking when the foci were near the bottom of the z-stack (see schematic in Fig. 4 D). We conclude that, when genomic loci can be tracked in 4D datasets, CNN-based denoising may introduce artifacts. Denoising becomes a more attractive solution when noise levels are very high (e.g. SNR < 1.5).

### Denoising single chromatin loci in yeasts cells

We tested if denoising using N2V would significantly alter the results of a single particle tracking experiment. Time lapse images of a 10 kb lacO/LacI-GFP array positioned 1.8 kb from centromere 15 and of spindle pole bodies labeled with Spc29-RFP (Fig. 5A) were background-subtracted and denoised using N2V. Low-dose benomyl treatment is known to restrict the motion of pericentric chromatin in yeast (Lawrimore *et al.*, 2015). The restriction of the motion of the foci in benomyl-treated cells was apparent in the original time lapse images as well as in the denoised time lapses. The motion of the spindle pole bodies was not restricted upon benomyl treatment and this result was recapitulated in the denoised time lapses (Fig. 5B). While denoising did introduce tracking errors (outliers in Fig. 5B-C) for a few loci, these did not not lead to statistically different results compared to the original images. Denoising maintained higher SNR over the time course (Fig. 5D), meaning that implementing N2V could allow longer observations.

**Figure 5.**
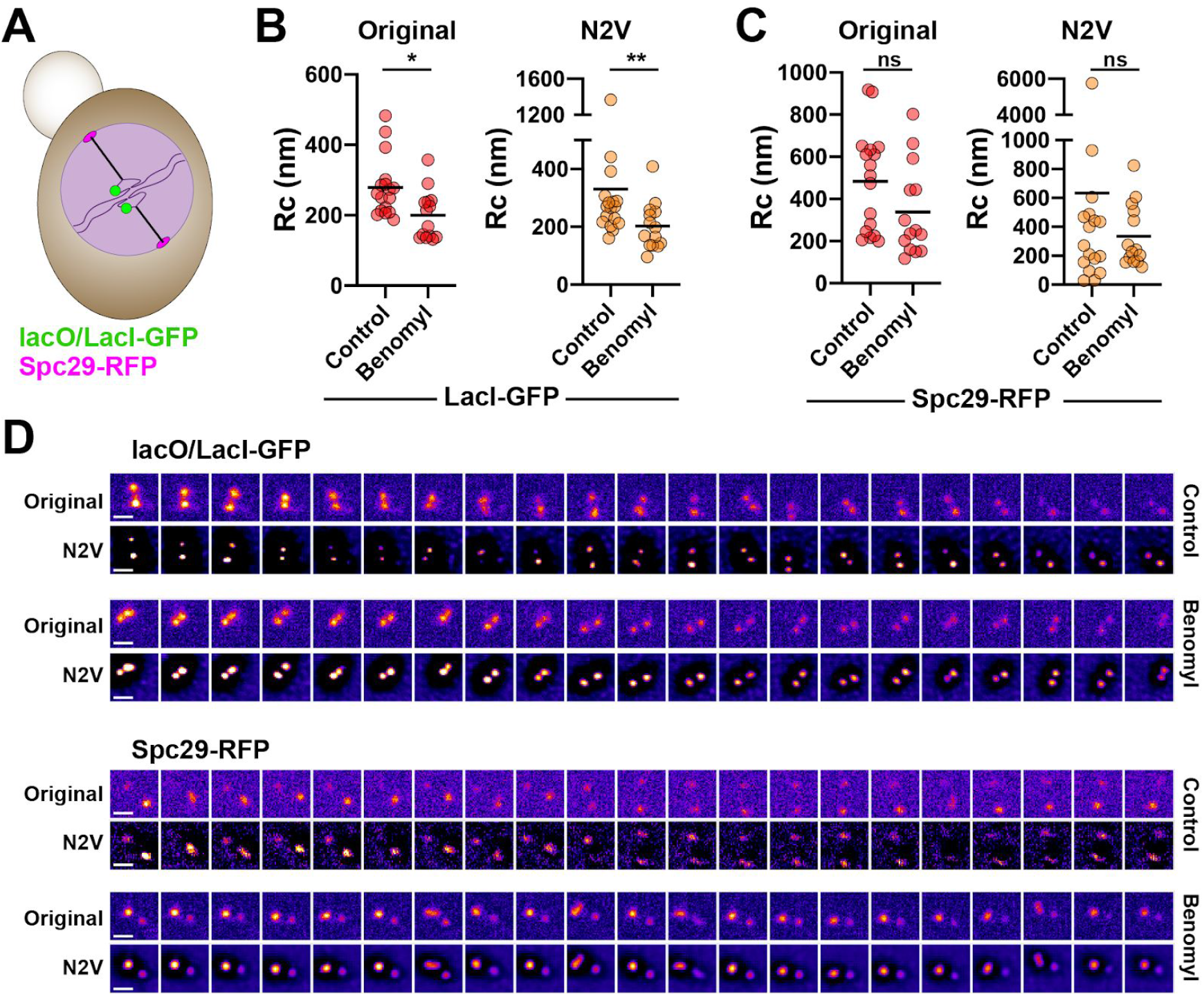
N2V denoising does not mask reduction of chromatin motion upon benomyl treatment. **A** Schematic of 10 kb lacO/LacI-GFP tandem repeat array located 1.8 kb from CEN15. The spindle body is labeled with Spc29-RFP. **B-C** Radius of confinement (Rc) for sister LacI-GFP signals (B) and Spc29-RFP (C) for background-subtracted images (original) and background-subtracted images denoised with N2V (N2V). *, *P* < 0.05; **, *P* < 0.01 (Mann Whitney test). Mean values are shown. **D** Representative time lapse montage of lacO/LacI-GFP sister foci signals and Spc29-RFP signals in original and N2V-denoised images of untreated or benomyl-treated, metaphase cells. Images are sum intensity projections with an interval of 30 seconds. Scale bars, 1 μm.

### A systematic approach to optimize the hyper-parameters of CNN networks

A non-trivial aspect of training CNN networks such as CARE is choosing the network’s hyper-parameters and deciding on the size of the training set. While some parameters such as the learning rate may be application-invariant and thus are generally kept as described by the authors (Weigert *et al.*, 2018), others such as the restoration grid size are more application-specific. Refining these hyper-parameters with a system may improve the restoration outcomes. Yet relying on trial-and-error is time-consuming. To rigorously identify hyper-parameters optimal for our type of data, we employed Bayesian optimization (Nogueira, 2014) (Fig. 6A). In this approach, the network’s parameter space is modeled using Gaussian processes. This optimization can find hyper-parameter values that lead to the best performance at the task at hand while minimizing the amount of evaluations. The approach is thus especially useful when the evaluation of the target function is expensive, as in our case in which an image restoration network needs to be trained and its predictions tracked at each evaluation step of the optimizer. We used three different objective functions for assessment: cumulative tracked motion in fixed cells, cumulative tracking error in live cells, and relative number of spots tracked. Fig. 6B shows the sets of optimal parameter values found by the Bayesian optimization process identified using these target functions. Compared to the default hyperparameter set of the CARE implementation used, the hyperparameters found through this optimization resulted in a model producing restorations with a 12.8% lower cumulative tracking error.

**Figure 6.**
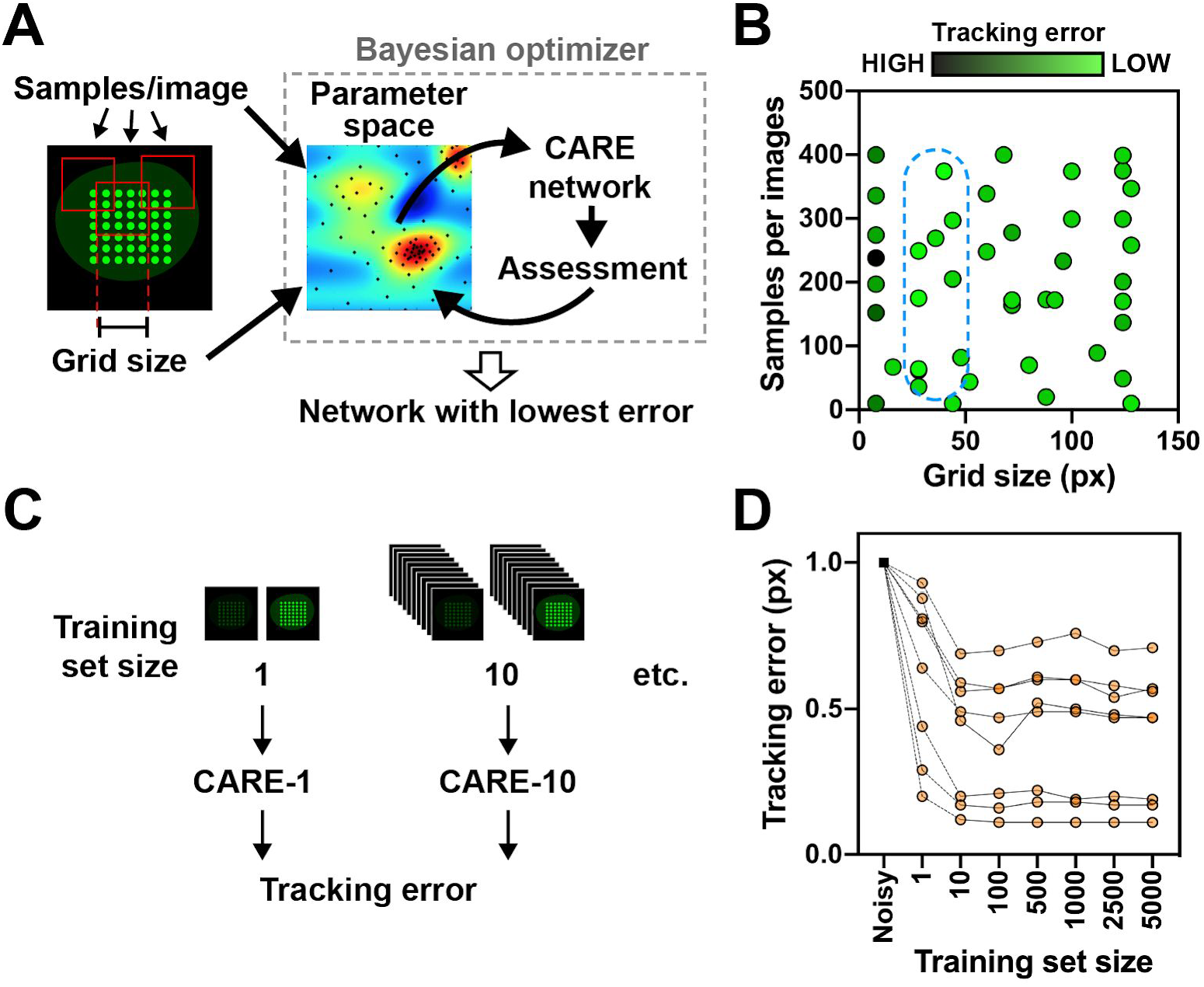
Optimization of CARE network hyperparameters. **A** Schematic of the approach to simultaneously assess multiple parameters using Baysian optimization. For chromatin microdomain tracking, the reward function minimized the cumulative tracking error. **B** Effect of the grid size and the number of training sub-samples on tracking accuracy. The graph shows that the optimizer sparsely explored the entire parameter space before focusing on the most promising areas (dotted line). Each dot on the graph represents the performance (tracking accuracy) of a CARE network, mapping noisy (10 ms exposure) images to GT (300 ms) images from interlaced live-cell movies. **C** Approach to determine optimal size of the training set, with CARE CNNs trained on different numbers of image pairs. **D** The tracking error, expressed relative to the original noisy images, is shown as a function of the training set size.

Producing large training sets for the supervised deep learning approach with matching noisy and clean images can be time consuming. To investigate the effect of training set size on the restoration outcome, we trained CARE networks with identical hyper-parameters on differently sized training sets, and quantified the change in tracking error (Fig. 6C-D). As expected, we found that models trained with more training data had a lower cumulative tracking error. A performance plateau was achieved with a surprisingly small number of training image pairs (10-100). This suggests that, at least in our experimental conditions, significant improvement in image quality and tracking precision can be achieved with a small dataset. We do not exclude the possibility that further improvements would have been achieved with a much larger training set, but considered that a very large set of matched noisy/GT image pairs is experimentally and computationally impractical.

## Conclusion

We evaluated the performance of content-aware deep learning methods for denoising microscopy image sequences, using particle tracking outcomes as an objective assessment. In contrast to conventional denoising approaches, CNN-based image restoration makes little or no assumptions on noise. As such, no pre-acquisition of camera-based noise features or calibrations is needed, which facilitates the implementation of these methods. By learning the noise pattern of a given image (or image pairs), both N2V and CARE remove noise regardless of its source (shot noise, dark noise, readout noise, etc), with the notable exception of structural noise for N2V. Overall, we find that the deep learning methods can restore biophysical information from very noisy datasets, highlighting their potential for quantitative microscopy. As expected, the supervised method (CARE) generally performed better than the unsupervised method (N2V), which led to structural noise features after restoration, in particular for single-nucleosome dataset. Our results with supervised learning are very encouraging because individual chromophores in single-molecule tracking experiments have a limited photon budget, necessitating minimal excitation light during image acquisition. CNN-based denoising in our analyses was particularly efficacious to restore single image planes, with mixed results obtained for image volumes. We also conclude that these methods are mostly relevant when noise levels are very high, and note the important caveat that, in contrast to classic denoising methods, content-aware deep learning approaches can fabricate biologically-irrelevant information, which needs to be carefully evaluated. We anticipate further improvements in deep learning denoising methods as the field rapidly expands, and propose that the approach presented here will be useful to rigorously assess their performances.

## MATERIALS AND METHODS

### Synthetic data to evaluate denoising performances for particle tracking in 2D

A synthetic model was designed by our lab where the number of particles, diffusion coefficient and noise can be defined for each movie. MSD was used to characterize particle motions. MSD is defined by the following equations:

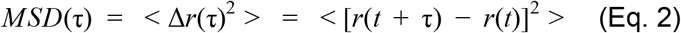

In two dimensions,

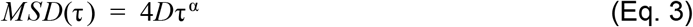

The term *D* in Eq. 3 is the diffusion coefficient of the particle and the exponent α is a unitless parameter which characterises the type of diffusion; α =1 for simple diffusion, and α =½ for a stretch of beads in a long-chain polymer (Osmanović and Rabin, 2017). In a Brownian motion model, the MSD is dependent on the size of the moving object as well as the mechanical and physical properties of the medium, as described in Eq. 4 (the Stokes-Einstein equation).

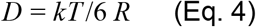

Here η is the viscosity of the medium, T is the temperature, *R* is the particle radius, and k is the Boltzmann constant. To match experimental observations, the pixel size of the synthetic data was set as 80 nm. The particle was simulated as a 2D Gaussian function with a radius of 60 nm. To avoid merging of multiple foci, the *D* value was set to 1.2 nm^2^/s. Movies including 49 particles lasting for 300 frames were simulated with a temporal interval of 1 s. The noise level of synthetic images was simulated as the Gaussian noise, with the standard derivation (σ) varies as σ = 0, 0.1, 0.2, 0.3, 0.4, 0.5, and 0.6. Synthetic movies without noise (σ = 0) were used as ground truth, and the coordinates of each particle were also used as ground truth for MSD curves. For restored images, the motion of the particle was tracked by a home-written algorithm (details below).

### Image generation and tracking of simulated genomic loci

We used ChromoShake (Lawrimore *et al.*, 2016) simulations of the budding yeast pericentric region to simulate yeast genomic loci. These simulations were converted into synthetic images using Microscope Simulator 2 (Quammen *et al.*, 2008). In Microscope Simulator 2, a custom point spread function was generated using the “Calculated Gibon-Lanni Widefield PSF” method for a 100x magnification, 1.49 numerical aperture objective. The custom point spread function was convolved with either 10 or 20 consecutive monomer units, representing 200 bp or 400 bp fluorescent reporter operator arrays, respectively. These were positioned at the apex of 64 different loops in the pericentric simulations by first removing the header from the ChromoShake outfile and converting the coordinates from meters to microns. The resulting text file was converted to a series of Microscope Simulator 2 model files. The model files were converted to TIFF stacks. The simulated TIFF stacks were concatenated into 4-dimensional hyperstacks (50×50 pixels, 5 z-planes, 200 nm z-step, 201 timepoints) using FIJI (Schindelin *et al.*, 2012). All simulations were converted into images containing either no noise (ground truth), or randomly distributed noise with a standard deviation of 5 AU. The gain of the signal of the noisy images was set to 100-fold, 1000-fold, and 10000-fold, to generate images with differing signal to noise ratios.

The simulated DNA loci were tracked in each 4D hyperstack using a custom MATLAB code that locates the brightest voxel in a z-stack, crops a 15×15×5 region surrounding the brightest voxel, and uses MATLAB’s lsqcurvefit function to fit a 3-dimensional Gaussian function to the cropped region. The center of the fitted Gaussian function was calculated for each timepoint to create a single track per timelapse. The ground truth track for each timelapse was calculated by taking the mean position of all the labeled masses directly from the simulation model XML file using a custom MATLAB program. The mean squared displacement and radius of confinement were calculated by custom MATLAB programs.

### Mammalian cell culture

U2OS osteosarcoma cells were cultured in DMEM supplemented with 10% fetal bovine serum (Sigma) at 37°C, 5% CO_2_. Cells were seeded in 35 mm glass-bottom dishes (MatTek) at 100,000 cells per dish and imaged 48h after seeding. U2OS cells stably expressing PAGFP-H2A (Bonin *et al.*, 2018) were used to track chromatin microdomains. For single nucleosome imaging, we generated U2OS cells stably expressing H2B fused to the HaloTag by transfection of the pBREBAC-H2BHalo plasmid (Addgene plasmid # 91564) using Lipofectamine 3000 (ThermoFisher) followed by clonal selection with geneticin. Before live cell imaging, H2B-HaloTag U2OS cells were incubated with 10 pM fluorescent JF 459 HaloTag ligand (Grimm *et al.*, 2015) for 1h, washed three times with PBS, and incubated in DMEM without phenol red for at least 30 min. This concentration of dye proved to be optimal for imaging and tracking. For fixed imaging, cells were imaged after fixation with formalin (Sigma # HT5011; 20 min).

### Chromatin microdomains tracking

Grids of photoactivated chromatin microdomains (7×7) were generated in U2OS PAGFP-H2A cells with a custom diffractive optical element module (Bonin *et al.*, 2018) inserted into the condenser arm of an inverted Olympus IX83 microscope. Cells were kept at 37°C in the custom-built enclosure of the microscope. Images were taken with a 60x oil lens (N.A. = 1.35) and an sCMOS camera (ORCA-Flash 4.0 v3; Hamamatsu Photonics; dark current *D_c_* = 0.2 electrons/pixel/s when cooled to −15°C), using the CellSens software. Images were registered using the StackReg plugin in ImageJ (Thévenaz *et al.*, 1998). Tracking of chromatin microdomains was done in MATLAB, as described (Bonin *et al.*, 2018).

### Single nucleosome tracking

Motion of single nucleosomes in live U2OS cells was tracked using a custom-built oblique light sheet microscope, based on the ASI’s rapid automated modular microscope (Applied Scientific Instrumentation). A solid-state laser (559 nm, 30 mW, Olympus) was focused by a tube lens to the side of the back pupil of an oil immersion objective (60x, N.A. = 1.2, Olympus), leading to a thin sheet of light that illuminated the cell. Fluorescent signals were collected by the same objective and further filtered by a multi-band emission filter (69013M, Chroma). Finally, the fluorescent signal was detected by an ORCA-Flash 4.0 sCMOS camera. Cells were maintained in a physiological environment using a live cell imaging chamber (INU-TIZ-F1; Tokai Hit). The microscope system and the time-course image acquisition were controlled by the open-source software MicroManager. Single nucleosome motions were analyzed with a custom single-molecule image analysis platform, smCellQuantifier. Specifically, the localization of each nucleosome locus was detected and fitted with a two-dimensional Gaussian function. Trajectories of single nucleosomes were established with a multitemporal association tracking algorithm (Shafique and Shah, 2005; Winter *et al.*, 2012). The MSD of each focus was calculated based on the trajectory profile, and the diffusion coefficient *D* as well as the anomalous coefficient α were calculated with equation 3.

### Tracking genomic loci in yeast

Budding yeast strain KBY8065 (Mat a CEN15(1.8)-GFP[10kb] ade2-1, his3-11, trp1-1, ura3-1, leu2-3,112, can1-100, LacINLSGFP:HIS3, lacO::URA3, Spc29RFP:Hyg) were grown in liquid yeast extract peptone dextrose at 24°C. Cells were imaged in liquid yeast complete medium at 24°C. Time lapse images were acquired on an Eclipse Ti wide-field inverted microscope (Nikon) with a 100x Apo TIRF 1.49 NA objective (Nikon) and a Clara charge-coupled device camera (Andor) using the Nikon NIS Elements imaging software. Time lapses were 10 min in duration with 30 s intervals. At each interval, a seven-step Z-stack of 400-nm step size was acquired in the GFP, RFP, and Trans channels.

Metaphase yeast cells (medium budded cells with two Spc29-RFP foci) were cropped by hand from the original time lapse images. The heterogeneous background was subtracted with the rolling-ball method (10 pixel radius) in FIJI. Both background-subtracted and denoised time lapses were automatically tracked using a custom MATLAB program. The motion of the two sister lacO/LacI-GFP foci and the two sister Spc29-RFP foci were the motion of one focus relative to the other as in Chacon et al. (Chacón *et al.*, 2014) and the radius of confinement was calculated by a custom MATLAB program.

### CNN training and Bayesian optimization of CNN hyperparameters

For synthetic bead data (2D), 100 pairs of noisy and GT images were used for training the CARE algorithm, using a patch size of 128×128 pixels. N2V networks were trained on images with high noise levels (*σ* = 0.5), using a patch size of 64×64 pixels.

For restoration of chromatin microdomain images, dedicated CARE networks were trained for each exposure time condition (1 ms, 3 ms, 10 ms) using pairs of cropped images taken from fixed cells. These training pairs were obtained by alternately imaging at the target exposure time (e.g. 3 ms) and an exposure time sufficient to achieve a high SNR (e.g. 300 ms) for 100 times, and subsequently cropping image stacks centered at the grid of photoactivated chromatin microdomains. The hyper-parameters for grid size (28×28 px) and samples per image (64) were chosen by applying the Bayesian optimization implementation by Nogueira (Nogueira, 2014), with the objective reward function of minimal cumulative tracking error. We calculated tracking error as pixels per microdomain per frame: we summed the magnitude of the difference between the motion vector tracked in the ground-truth and the motion vector tracked in the denoised image for each frame-delta across all spots, and then divide by the number of frame-deltas and spots. N2V networks for use with the chromatin microdomain images were trained on the cropped target image stacks directly, with one network trained for each image stack. We evaluated performance on training N2V networks on the full images (2048×2048 px), but found no improvement. Rather training on the full images significantly increased training time. For both CARE and N2V we used a 90%-10% train-validation split.

To restore single nucleosome images, a CARE network was trained using 100 pairs of fixed cell images, captured at 1 s and 10 ms. N2V was trained using stacks of fixed images (100, 10 ms exposure). We optimized patch size (128×128 pixels for CARE and 64×64 for N2V) prior to training the networks. Since we had high-quality fixed cell images (1 s exposure) as the GT, we were able to decide on the most accurate network.

For yeast genomic loci and spindle pole bodies, background-subtracted images were processed using 3D N2V to generate denoised timelapses. For both simulated genomic loci and spindle pole body timelapses, one N2V network was trained for each observation, with a patch size of 32, 32, and 4 px for X, Y, and Z, respectively.

### Statistical analyses

Statistical analyses were done using GraphPad Prism 8. The D’Agostino & Pearson omnibus normality test was used to test for normality. Nonparametric tests were used if the data did not pass the normality test (at alpha = 0.05). Statistical tests are indicated in the figure legends. *P*-values ≤ 0.05 were considered significant. All statistical tests were two-sided.

## ACKNOWLEDGEMENTS

We thank Dr. Jacques Neefjes (the Netherland Cancer Institute) for providing the PAGFP-H2A DNA construct and Dr. Zhe Liu (Janelia Research Campus, HHMI) for sharing the LZ10 PBREBAC-H2BHalo DNA plasmid. We are grateful to Dr. George Holzwarth for his comments on the manuscripts, for his contribution in developing MATLAB code, and for productive discussions. We also like to acknowledge Clayton Seitz and Dr. Hua Lin for contributing the single molecule tracking software and the synthetic image generator for this project. This work was funded by the National Cancer Institute (U01CA214282 and P30CA012197 to the Wake Forest Baptist Comprehensive Cancer Center).

### Box 1

**Box 1:**
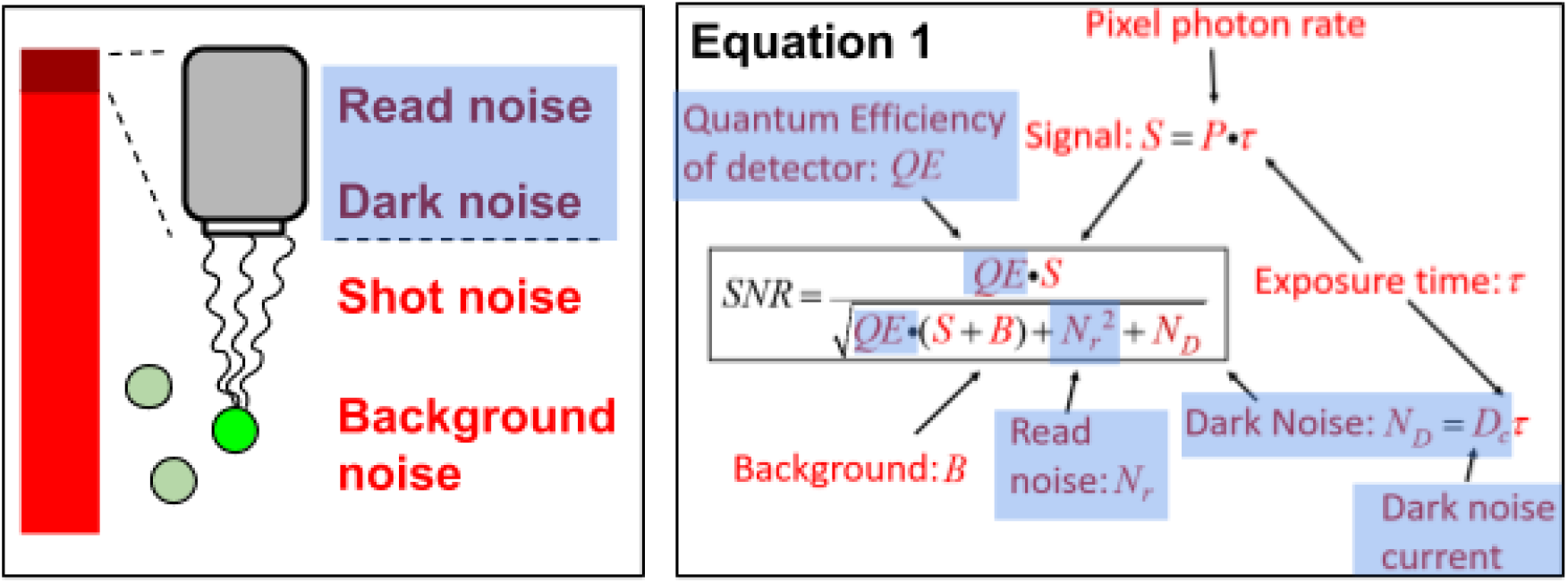
Different sources of noise in (cellular) imaging. In the schematic, noise dependent on the camera is color-coded bordeaux on blue background, whereas noise dependent on the sample is coded in red. The bar indicates relative proportions of sample- and camera-related noise in typical imaging conditions. In Eguation 1, *S* = *Pt* Is the signal [P is the incident photon flux (photons/pixel/second) and *t* is the exposure time (seconds)], QE represents the camera quantum efficiency (# electrons generated/incident photon), *B* is the background (same units as *S*), *D_c_* is the dark current value (electrons/pixel/second), and *N_r_* represents camera read noise (electrons rms/pixel).

## SUPPLEMENTARY MATERIAL

**Figure S1.**
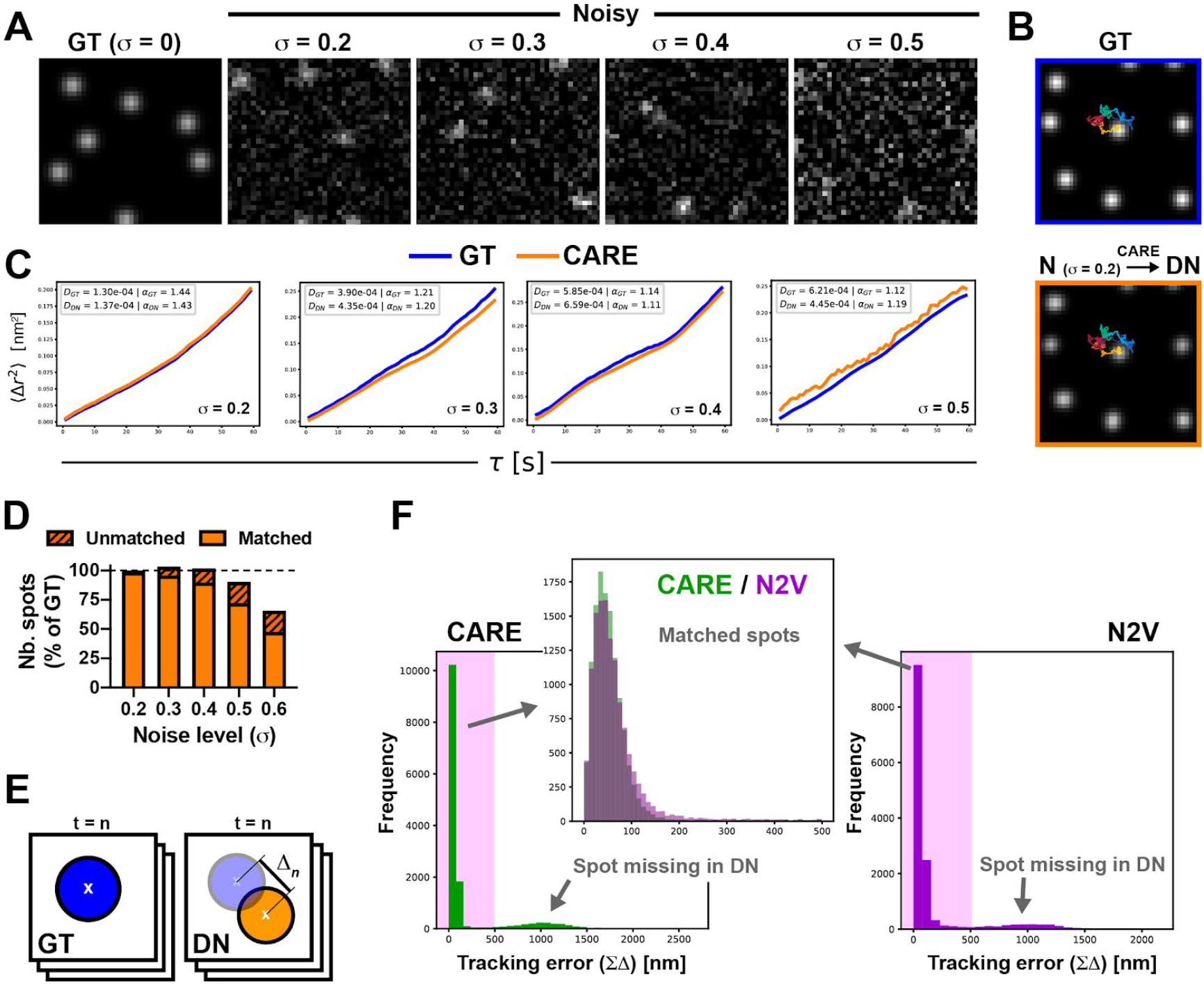
Evaluation of CNN-based denoising with synthetic data of diffusing beads (related to Fig. 1). **A** Illustration of synthetic bead images with different levels of noise (*σ*). **B** Trajectories of a bead in ground truth (GT) and denoised (DN) movies. The noisy movie used for denoising had a noise level of *σ* = 0.2. **C** MSDs calculated from single diffusing beads in GT and denoised movies. Denoised movies were generated from image sequences with different noise levels. **D** Proportion of bead spots identified in denoised movies (CARE), either matching or not matching a bead in the GT movie. **E** Approach to evaluate tracking accuracy. For all beads in GT movies, spot centers were determined in each image of the sequence (300). The distances to the closest spot center in the corresponding denoised movie (Δ, tracking error) were calculated. The Δ for all beads and all frames were summed. **F** Histograms of tracking errors for movies with a noise level of *σ* = 0.5, denoised with CARE or N2V.

**Figure S2.**
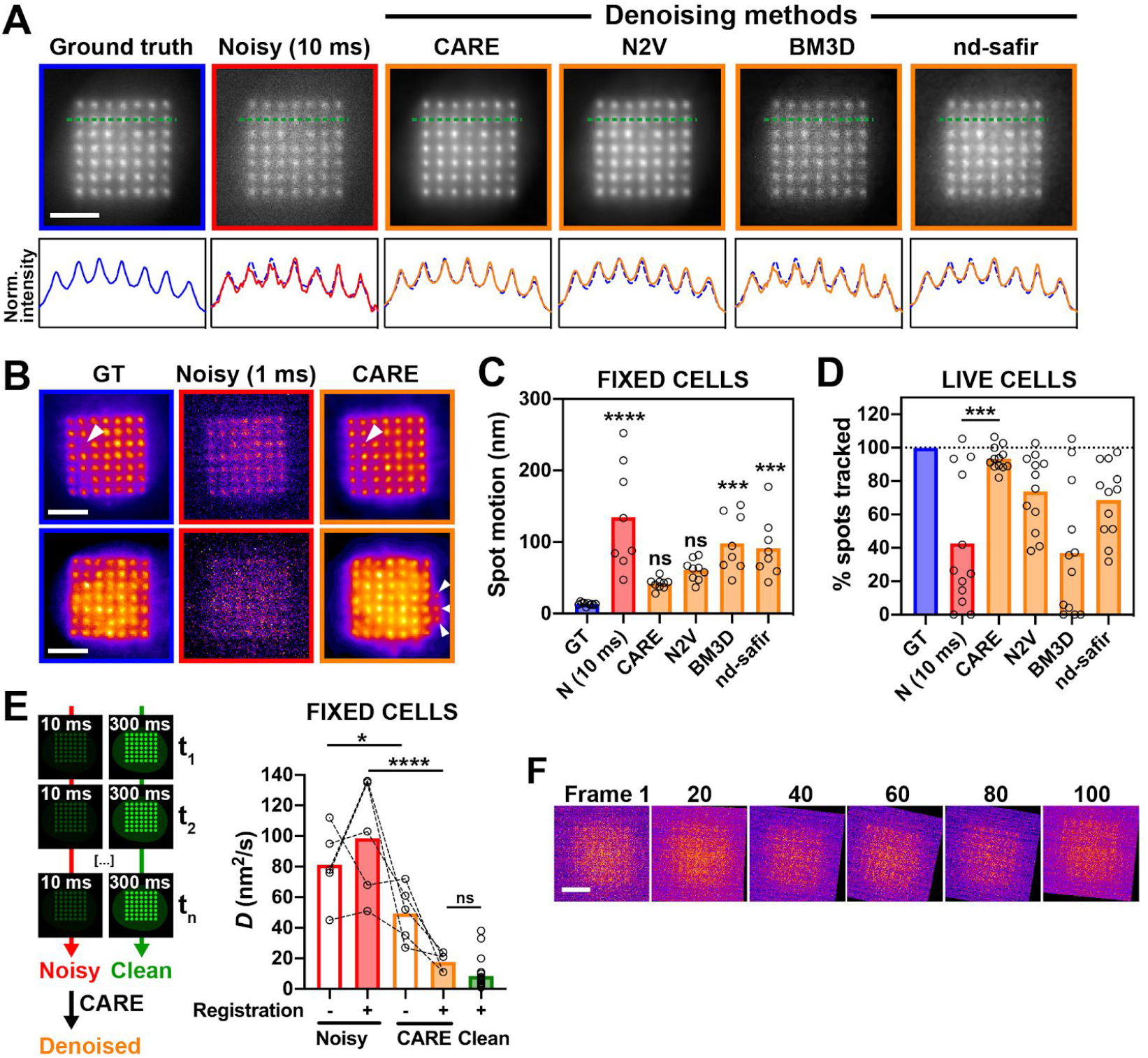
Image restoration to quantify the dynamics of chromatin microdomains in U2OS cells expressing PAGFP-H2A (related to Fig. 2). **A** Denoising of movies with low noise levels (10 ms exposure time) using different software, as indicated. Intensity profiles along the dotted line are shown. **B** Denoising of movies with very low exposure time (1 ms) with CARE CNN. Arrowheads highlight hallucinated spots. **C** Spot motions in fixed cells (10 ms). **D** Percentages of photoactivated spots that could be tracked in noisy movies (10 ms) or after denoising with the different algorithms. For each cell, values are normalized to the number of spots tracked in the GT. **E** Chromatin diffusion coefficient (*D*) derived from noisy (10 ms exposures), CARE-denoised, or ‘clean’ (300 ms exposure) image sequences taken from fixed cells. *D* values were calculated before and after registration of the time-lapses. Each dot in the graphs represents the average chromatin diffusion for a cell. *, *P* < 0.05; ***, *P* < 0.0005; ****, *P* < 0.0001; ns, not significant (ANOVA and Tukey’s test). Comparisons in C are relative to GT. **F** Illustration of image registration failure for a short exposure image sequence. Scale bars, 5 μm.

**Figure S3.**
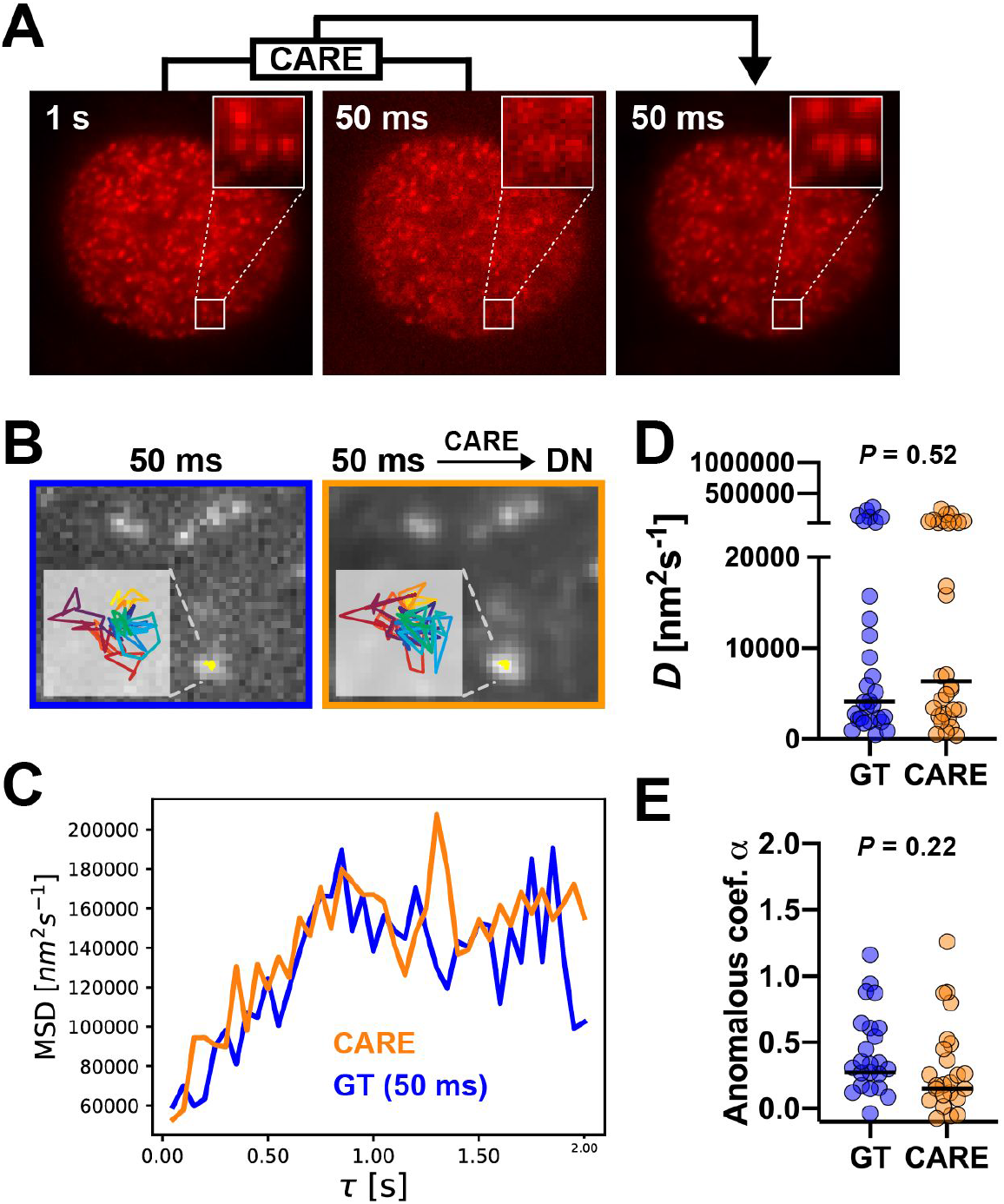
Image restoration to quantify motions of single nucleosomes (related to Fig. 3). **A** Images of U2OS H2B-HaloTag cell nuclei taken with long (1 s) or shorter (50 ms) exposure times. The CARE algorithm was used for denoising. **B** Single-nucleosome trajectories tracked in 50 ms exposure movies and corresponding movies after CARE denoising. **C** Average MSD traces of single nucleosomes from (B). **D** Diffusion values (*D*) and **F** anomalous coefficients derived from single-nucleosome MSD curves. Statistical comparisons with Mann-Whitney test. Median values are shown.

**Table S1.** Mean Radius of confinement values from noised and denoised image. Data is the same as in Figure 4D. *P*-values were calculated using a Kruskal-Wallis and Dunn’s multiple comparison test.

|  | Noised Image<br>Mean Rc<br>(nm) | Denoise Image<br>Mean Rc<br>(nm) | Nosied vs<br>Denoised<br>*P*-value | Noised vs<br>Ground Truth<br>*P*-value | Denoised vs<br>Ground Truth<br>*P*-value |
| --- | --- | --- | --- | --- | --- |
| 200 bp array,<br>100-fold gain | 1,148 | 216 | < 0.0001 | < 0.0001 | < 0.0001 |
| 200 bp array,<br>1000-fold<br>gain | 149 | 220 | 0.0092 | < 0.0001 | < 0.0001 |
| 200 bp array,<br>10000-fold<br>gain | 110 | 120 | 0.81 | > 0.99 | 0.50 |
| 400 bp array,<br>100-fold gain | 173 | 214 | 0.32 | < 0.0001 | < 0.0001 |
| 400 bp array,<br>1000-fold<br>gain | 120 | 199 | < 0.0001 | 0.052 | < 0.0001 |
| 400 bp array,<br>10000-fold<br>gain | 103 | 127 | 0.0003 | 0.0001 | 0.0001 |
| 200 bp array,<br>Ground truth | 109 | NA | NA | NA | NA |
| 400 bp array,<br>ground truth | 102 | NA | NA | NA | NA |

## Notes

### Competing Interest Statement

The authors have declared no competing interest.

